# Rapid, precise, and reliable measurement of delay discounting using a Bayesian learning algorithm

**DOI:** 10.1101/567412

**Authors:** Woo-Young Ahn, Hairong Gu, Yitong Shen, Nathaniel Haines, Hunter A. Hahn, Julie E. Teater, Jay I. Myung, Mark A. Pitt

## Abstract

Machine learning has the potential to facilitate the development of computational methods that improve the measurement of cognitive and mental functioning. In three populations (college students, patients with a substance use disorder, and Amazon Mechanical Turk workers), we evaluated one such method, Bayesian adaptive design optimization (ADO), in the area of delay discounting by comparing its test-retest reliability, precision, and efficiency with that of a conventional staircase method. In all three populations tested, the results showed that ADO led to 0.95 or higher test-retest reliability of the discounting rate within 10-20 trials (under 1-2 minutes of testing), captured approximately 10% more variance in test-retest reliability, was 3-5 times more precise, and was 3-8 times more efficient than the staircase method. The ADO methodology provides efficient and precise protocols for measuring individual differences in delay discounting.

---

Delay discounting, one dimension of impulsivity^1^, assesses how individuals make trade-offs between small but immediately available rewards versus large but delayed rewards. Delay discounting is broadly linked to normative cognitive and behavioral processes, such as financial decision making^2^, social decision making^3^, and personality^4^, among others. Also, individual differences in delay discounting are associated with several cognitive capacities, including working memory^5^, intelligence^6^, and top-down regulation of impulse control mediated by the prefrontal cortex^7,8^.

Delay discounting is a strong candidate endophenotype for a wide range of maladaptive behaviors, including addictive disorders^9,10^ and health risk behaviors^for a review see, 11^. Studies of test-retest reliability of delay discounting demonstrate reliability both for adolescents^12^ and adults^13^, and genetics studies indicate that delay discounting may be a heritable trait^9^. As such, delay discounting has received attention in the developing field of precision medicine in mental health as a potentially rapid and reliable (bio)marker of individual differences relevant for treatment outcomes^14-16^. The construct validity of delay discounting has been demonstrated in numerous studies. For example, the delay discounting task is widely used to assess (altered) temporal impulsivity of various psychiatric disorders, including patients with substance use disorders^e.g., 11^, schizophrenia^13,14^, and bipolar disorder^14^. Therefore, improved assessment of delay discounting may be beneficial to many fields, including psychology, neuroscience, medicine^15^, and economics.

To link decision making tasks to mental functioning is a formidable challenge that requires simultaneously achieving multiple measurement goals. We focus on three aspects of measurement: reliability, precision, and efficiency. Reliable measurement of latent neurocognitive constructs or biological processes, such as impulsivity, reward sensitivity, or learning rate, is difficult. Recent advancements in neuroscience and computational psychiatry^16,17^ provide novel frameworks, cognitive tasks, and latent constructs that allow us to investigate the neurocognitive mechanisms underlying psychiatric conditions; however, their reliabilities have not been rigorously tested or are not yet acceptable^18^. A recent large-scale study suggests that the test-retest reliabilities of cognitive tasks are only modest^19^. Even if a test is reliable across time, confidence in the behavioral measure will depend on the precision of each measurement made. To our knowledge, few studies have rigorously tested the precision of measures from a neurocognitive test. Lastly, cognitive tasks developed in research laboratories are not always efficient, often taking 10-20 minutes or more to administer. With lengthy and relatively demanding tasks, participants (especially clinical populations) can easily fatigue or be distracted^20^, which can increase measurement error due to inconsistent responding. A by-product of low task efficiency is that the amount of data (e.g., number of participants) typically available for big data approaches to studying psychiatry is smaller than in other fields.

Several methods using fixed sets of choices currently exist to assess delay discounting, such as self-report questionnaires^21^ and computer-based tasks. The monetary choice questionnaire^22^ is an example, which contains 27 multiple choice questions and exhibits 5- and 57-week test-retest reliability of *r*’s within .7-.8 in college students. Other delay discounting tasks use some form of an adjustment procedure in which individuals’ previous responses are used to determine subsequent trials based on heuristic rules for increasing or decreasing the values presented. Such methods often adjust the amount of the immediate reward^22^, with the goal of significantly reducing the number of trials required to identify discounting rates. However, many adjustment procedures still require dozens of trials to produce reliable results; a notable exception is the 5-trial delay discounting task, which uses an adjusting-amount method to produce meaningful measures of delay discounting in as few as five trials^23^. While it is difficult to imagine a more efficient task, its precision and test-retest reliability have not been rigorously evaluated. More generally, although the use of heuristic rules to inform stimulus selection can be an effective initial approach to improving experiment efficiency, such rules often lack a theoretical (quantitative) framework that can justify the rule.

Bayesian adaptive testing is a promising machine-learning method that can address the aforementioned challenges in efficiency, precision, and reliability to improve the study of individual difference in decision making using computer-based tasks^24,25^. It originates from optimal experimental design in statistics^26^ and from active learning in machine learning^27^. Adaptive design optimization (ADO; **Figure 1**), an implementation of Bayesian adaptive testing, is a general-purpose computational algorithm for conducting adaptive experiments to achieve the experimental objective with the fewest possible number of observations. The ADO algorithm is formulated on the basis of Bayesian statistics and information theory, and works by using a formal cognitive model to guide stimulus selection in an optimal and efficient manner. Stimulus values in an ADO-based experiment are not predetermined or fixed, but instead are computed on the fly adaptively from trial to trial. That is, the stimulus to present on each trial is obtained by judiciously combining participant responses from earlier trials with the current knowledge about the model’s parameters so as to be most informative with respect to the specific objective. The chosen stimulus is optimal in that it is expected to reduce the greatest amount of uncertainty about the unknown parameters in an information theoretic sense. Accordingly, there would be no “wasted”, uninformative trials; evidence can therefore accumulate rapidly, making the data collection highly efficient.

**Figure 1.**
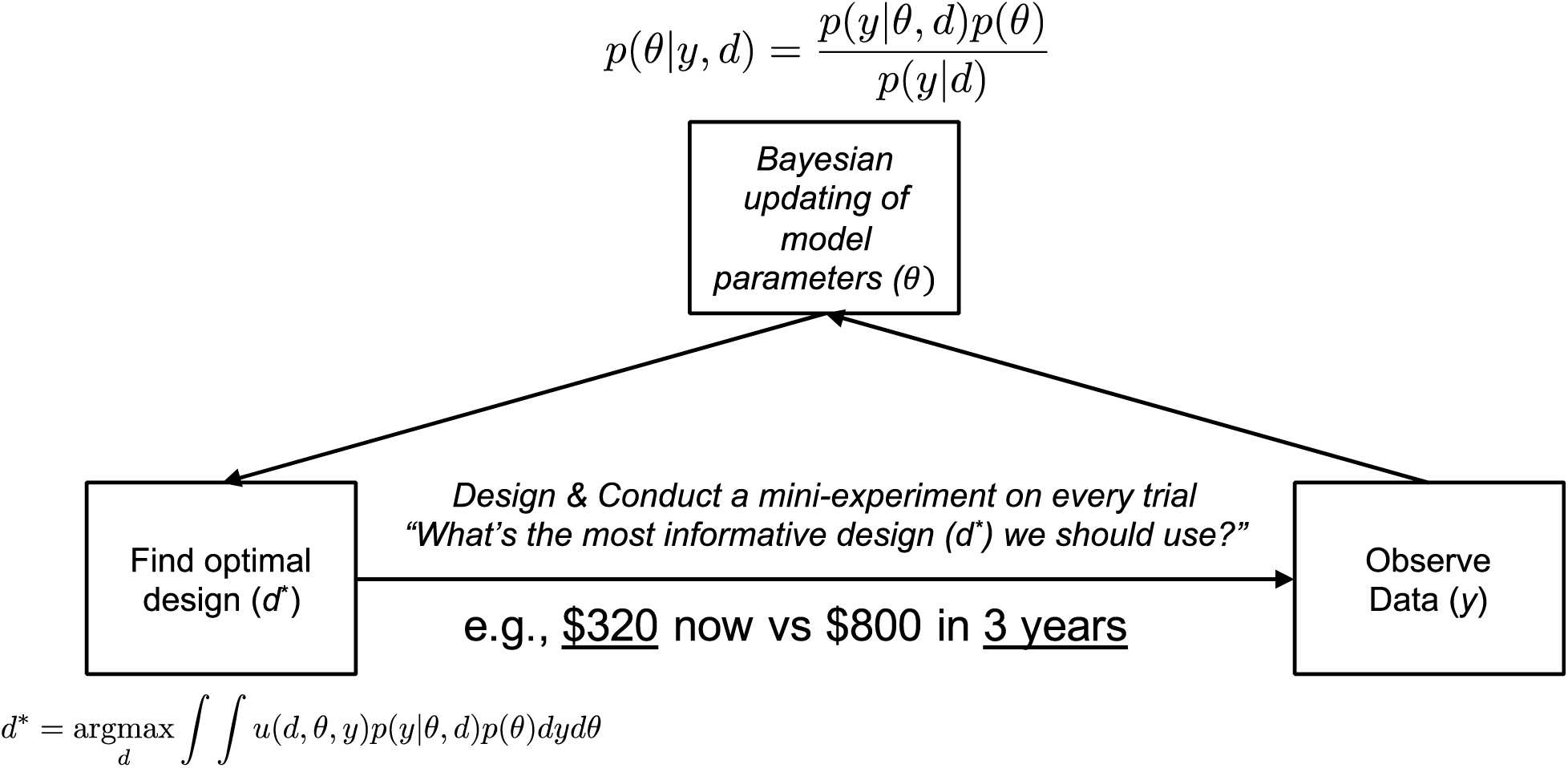
Schematic illustration of adaptive design optimization (ADO) in the area of delay discounting. Unlike the traditional experimental method, ADO aims to find the optimal design that extracts the maximum information about a participant’s model parameters on each trial. In other words, ADO identifies the most informative or optimal design (*d*^*\**^) using the participant’s previous choices (*y*), the mathematical model of choice behavior, and the participant’s model parameters (*θ*). In our delay discounting experiment with ADO, *y* would be 0 (choosing smaller and sooner reward) or 1 (larger and later reward), the mathematical model would be the hyperbolic function (see **Methods**), *θ* would be *k* (discounting rate) and *β* (inverse temperature), and *d\** would be the experimental design (a later delay and a sooner reward, which are underlined in the figure) that maximizes the integral of the local utility function, *u*(*d, θ, y*), which is based on the mutual information between model parameters (*θ*) and outcome random variable conditional upon design (*y*|*d*). For more mathematical details of the ADO method, see^24,25^.

ADO and its variants have recently been applied across disciplines to improve the efficiency and informativeness of data collection (cognitive psychology^28,29^, vision^30,31^, psychiatry^32^, neuroscience^33,34^, clinical drug trials^35^, and systems biology^36^).

Here, we demonstrate the successful application of adaptive design optimization (ADO) to improving measurement in the delay discounting task. We show that in three different populations (college students, patients with substance use disorders, and Amazon Mechanical Turk workers), ADO leads to rapid, precise, and reliable estimates of the delay discounting rate (*k*) with the hyperbolic function. Test-retest reliability of *k* reached up to 0.95 or higher within 10-20 trials (under 1-2 minutes of testing, including practice trials) with at least three times greater precision and efficiency than a staircase method that updates the immediate reward on each trial^37^. It is worth noting here that while in the present study we used ADO for efficiently estimating model parameters with a single (hyperbolic) model, ADO can also be, and has been, used to discriminate among a set of multiple models for the purpose of model selection^29,38-40^.

## Results

In Experiment 1, we recruited college students (N=58) to evaluate test-retest reliability (TRR) of the ADO and staircase (SC) methods of the delay discounting task over a period of approximately one month, a span of time over which one might want to measure changes in impulsivity. Previous studies have typically used 1 week^e.g., 41^, 2 weeks^42^, or 3-6 months^43^ between test sessions to evaluate TRR. Students visited the lab twice. In each visit they completed two ADO and two SC sessions, allowing us to measure TRR within each visit and between the two visits. In each task (or session), students made 42 choices about hypothetical scenarios involving a larger but later reward versus a smaller but sooner reward. We examined TRR using concordance correlation coefficients^44^ within each visit and between the two visits, using the discounting rate *k* of the hyperbolic function as the outcome measure (see **Methods**).

Past work customized the SC method to yield very good TRR^11^. Consistent with previous studies, in visits 1 and 2 of Experiment 1, within-visit TRR were 0.903 (visit 1) and 0.946 (visit 2), respectively. Nevertheless, ADO improved on this performance of SC, yielding values of 0.961 and 0.982, an increase of 10.8% (visit 1) and 6.9% (visit 2) in terms of the amount of variance accounted for (**Figure S1; Figures S2-3** shows the results for all participants including the outliers with ADO and SC, respectively – see **Methods** for the criteria for outliers). We found that TRR was higher at visit 2 than at visit 1. This was true of ADO and SC, and thus is likely indicative of participants learning the task and better adapting themselves to it in the second session (‘the practice effect’). Where ADO excels over the SC method is in efficiency and precision. We measured the efficiency of the method by calculating how many trials are required to achieve 0.9 TRR of *k*, which was assessed cumulatively at each trial (**Figure 2**). We measured precision using within-subject variability, quantifying it as the standard deviation (SD) of an individual parameter posterior distribution. With ADO, while we should evaluate its performance with fewer than several trials rather cautiously, we achieved over 0.9 TRR within 7 trials at visit 1. At visit 2, TRR exceeded 0.9 within 6 trials. With the SC method, TRR failed to reach 0.9 even at the end of the experiment (42 trials) at visit 1, and reached 0.9 only after 39 trials at visit 2. Although 0.9 TRR is an arbitrary threshold, it was chosen because it is stringent. Conclusions do not change qualitatively even if a more conservative or liberal threshold is used. For example, with the threshold of 0.8, the TRR with ADO reached the threshold within 2 (visit 1) and 3 (visit 2) trials in Experiment 1. On the other hand, the TRR with SC reached the threshold within 24 (visit 1) and 33 (visit 2) trials. If we set the threshold to 0.95, the TRR with ADO reached the threshold within 24 (visit 1) and 8 (visit 2) trials. The SC failed to reach the threshold even after 42 trials. Furthermore, ADO yielded approximately 3-5 times more precise estimates of discounting rate as measured by the smaller standard deviation of the posterior distribution of the parameter, *k* (ADO visit 1: 0.122, visit 2: 0.098; SC visit 1: 0.413, visit 2: 0.537; **Figure S4**).

**Figure 2.**
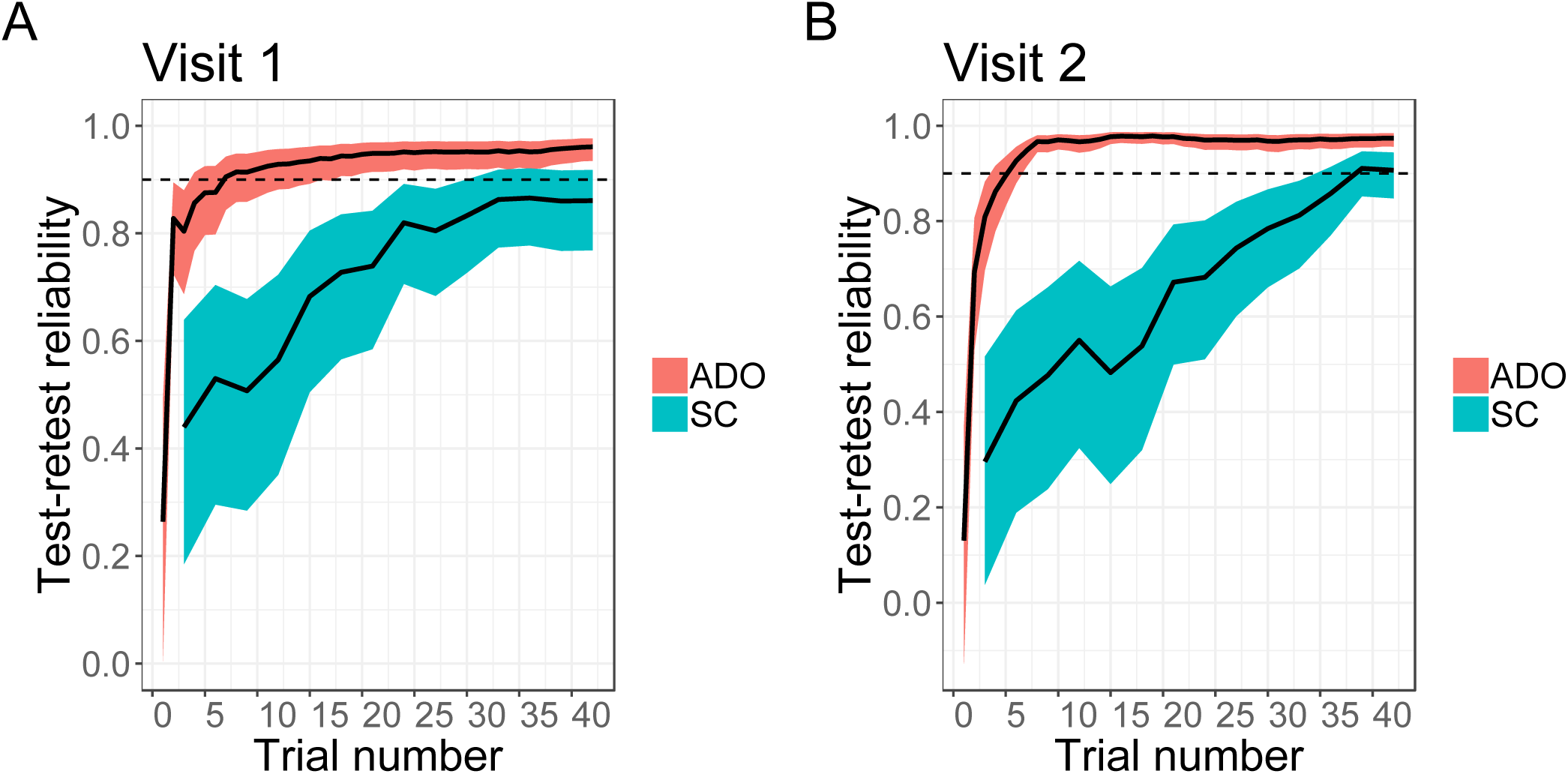
Comparison of ADO and Staircase (SC) within-visit test-retest reliability of temporal discounting rates when assessed cumulatively in each trial (ADO) or every third trial (SC) (Experiment 1, college students) at each of the two visits. Two visits were separated by approximately one month. In each visit, a participant completed two ADO sessions and two SC sessions (within-visit test-retest reliability). Test-retest reliability was assessed cumulatively in each trial. Shaded regions represent ±1 standard deviation across participants.

ADO also showed superior performance when examined across visits separated by one month (**Figure S5**). All four TRR measures across the two visits converged at around 0.8 within 10 trials and were highly consistent with each other. In contrast, with the SC method, the trajectories of the four measures were much more variable and asymptote, if at all, below 0.8 towards the end of the experiment.

In addition, we investigated if participants with low inverse temperature rate (*β*) at visit 1 show greater discrepancy in k estimates across visits 1 and 2. With ADO, we found small but significant negative correlations (r < −0.270, p < 0.04). This result suggests that participants who are less deterministic in their choices at visit 1 tend to show a larger discrepancy in *k* across the two visits. However, with the SC method, none of the correlations were significant (p > 0.11). We believe this outcome is due to the fact that many participants’ inverse temperature rate with the SC method are less distributed and clustered near zero in comparison to those estimated with ADO (see **Figure S2** for ADO estimates and **Figure S3** for SC estimates). Overall, the results of Experiment 1 show that ADO leads to rapid, reliable, and precise measures of discounting rate.

In Experiment 2, we recruited 35 patients meeting the Diagnostic and Statistical Manual of Mental Disorders (5^th^ ed. DSM-5) criteria for a substance use disorder (SUD) to assess the performance of ADO in a clinical population. The experimental design was the same as in Experiment 1 except that there was only a single visit. **Figure 3A** show that even in the patient population, ADO still led to rapid, reliable, and precise estimates of discounting rates, again outperforming the SC method. With ADO, maximum TRR was 0.973 within approximately 15 trials. Consistent with the results of Experiment 1, the SC method led to a smaller maximum TRR (0.892) and it took approximately 25 trials to reach this maximum. Precision of the parameter estimate was five times higher when using ADO than the SC method (0.073 vs. 0.371). **Figures S6** shows the results for all participants including the outliers in Experiment 2. Initially we set the upper bound of *k* to 0.1 assuming it would be sufficiently large for patients, but found that some patients’ *k* values reached ceiling. After recruiting 15 patients, we set the upper bound of *k* to 1 and no patient’s *k* reached ceiling. **Figure S7** suggests that the results largely remain the same whether we exclude those patients whose k reached ceiling or not.

**Figure 3.**
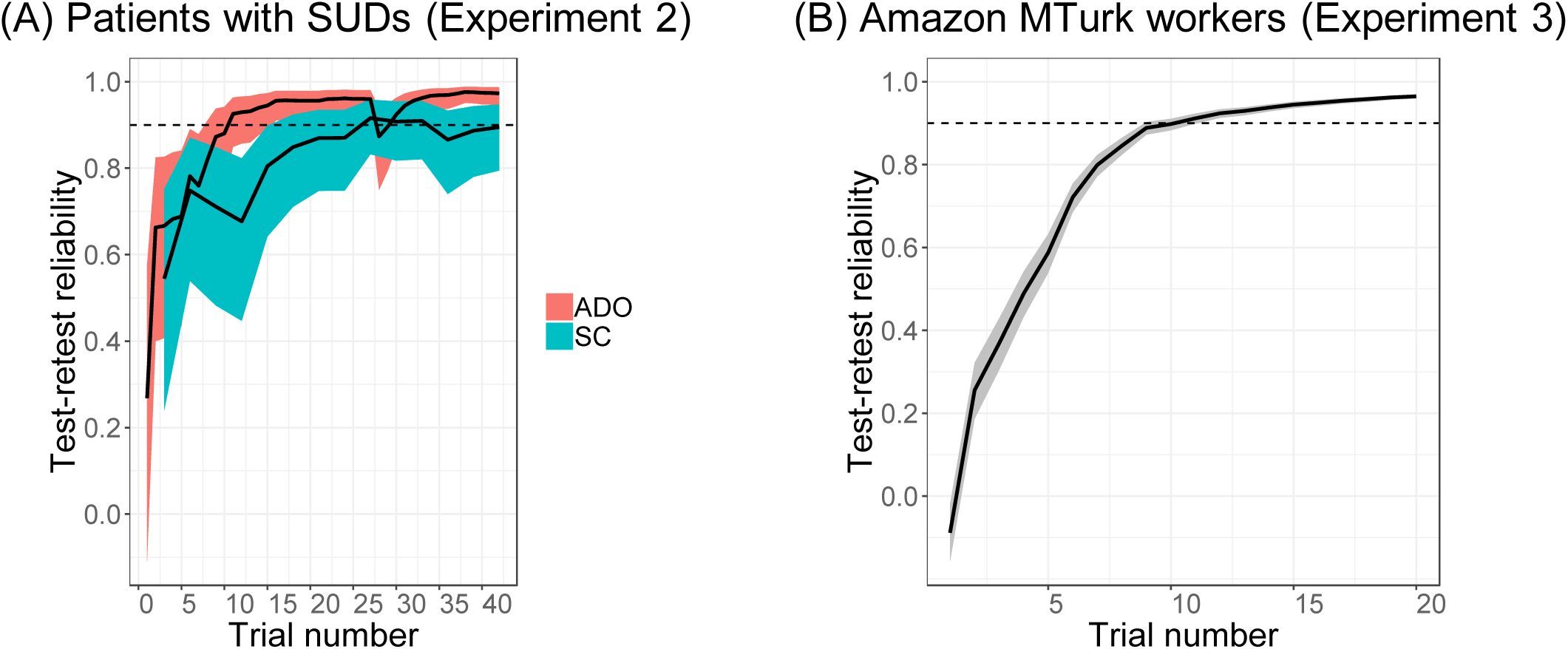
Reliability and efficiency of the ADO method in Experiments 2 and 3 (A) Comparison of ADO and Staircase (SC) test-retest reliability of temporal discounting rates when assessed cumulatively in each trial (ADO) or every third trial (SC) (Experiment 2, patients with substance use disorders (SUDs)) (B) Test efficiency as measured by the cumulative test-retest reliability across trials (Experiment 3, Amazon MTurk workers). Dashed line = 0.9 test-retest reliability. Unlike Experiments 1 and 2, only ADO sessions were administered and each session consisted of 20 trials in Experiment 3. Shaded regions represent ±1 standard deviation across participants.

In Experiment 3, we evaluated the durability of the ADO method, assessing it in a less controlled environment than the preceding experiments and with a larger and broader sample of the population, (808 Amazon Mechanical Turk workers). Each participant completed two ADO sessions, each of which consisted of 20 trials, which was estimated from Experiments 1 and 2 to be sufficient. In Experiment 3, ADO again led to an excellent maximum TRR (0.965), greater than 0.9 TRR within 11 trials as shown in **Figure 3B. Figure S8** shows the results for all participants, including outliers.

**Table 1** summarizes all results across the three experiments. Comparison of the two methods clearly shows that ADO is (1) more reliable (capturing approximately 7-11% more variance in TRR), (2) approximately 3-5 times more precise (smaller SD of individual parameter estimates), and (3) approximately 3-8 times more efficient (fewer number of trials required to reach 0.9 TRR). As might be expected, when tested in a less controlled environment (Experiment 3), precision suffers (0.339), being only slightly less than that found with the SC method (0.371), while reliability and efficiency hardly change.

**Table 1.**
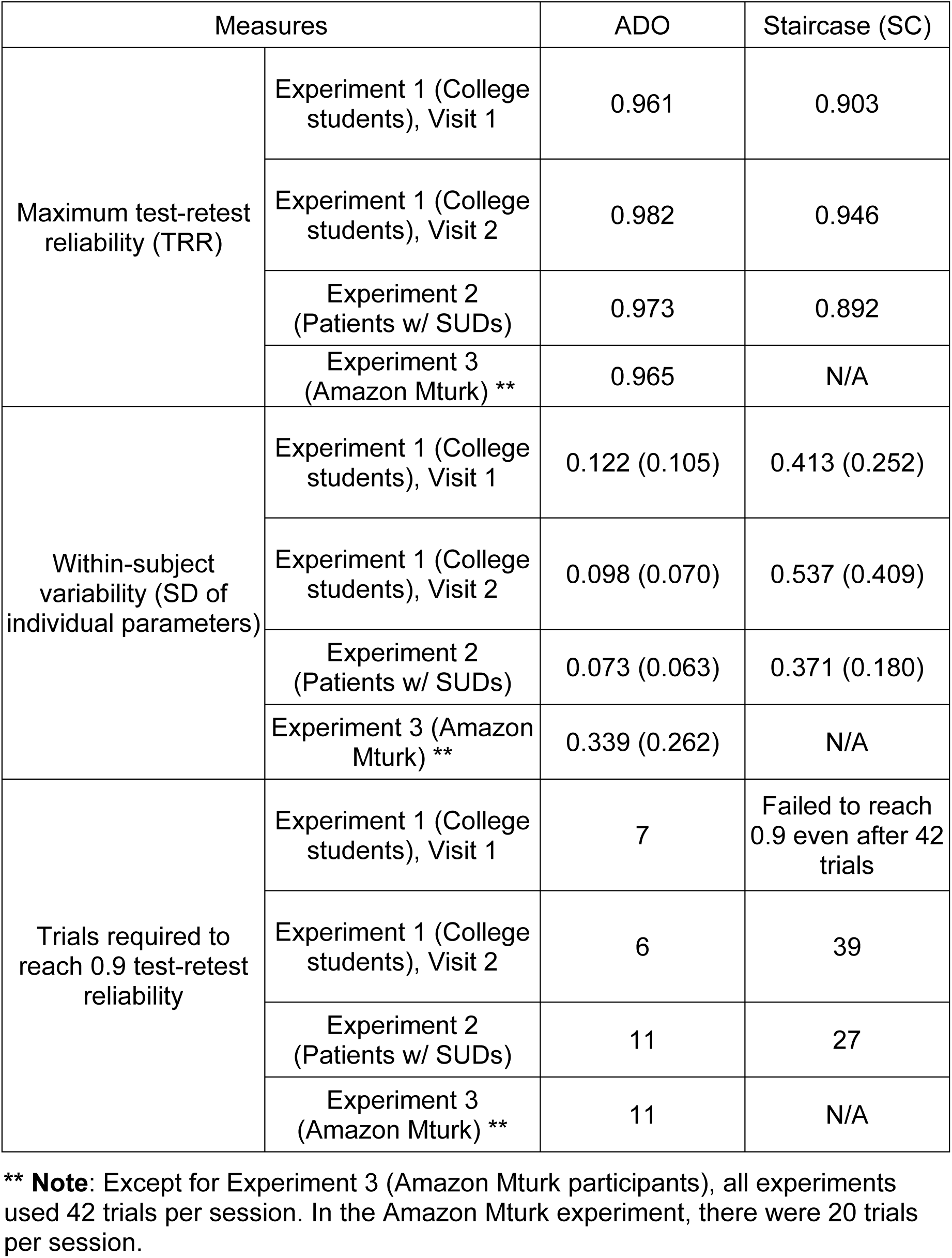
Comparison of ADO and Staircase (SC) methods in their reliability, precision, and efficiency (see **Methods** for their definitions) of estimating temporal discounting rates (*k*).

| Measures |  | ADO | Staircase (SC) |
| --- | --- | --- | --- |
| Maximum test-retest reliability (TRR) | Experiment 1 (College students), Visit 1 | 0.961 | 0.903 |
|  | Experiment 1 (College students), Visit 2 | 0.982 | 0.946 |
|  | Experiment 2 (Patients w/ SUDs) | 0.973 | 0.892 |
|  | Experiment 3 (Amazon Mturk) ** | 0.965 | N/A |
| Within-subject variability (SD of individual parameters) | Experiment 1 (College students), Visit 1 | 0.122 (0.105) | 0.413 (0.252) |
|  | Experiment 1 (College students), Visit 2 | 0.098 (0.070) | 0.537 (0.409) |
|  | Experiment 2 (Patients w/ SUDs) | 0.073 (0.063) | 0.371 (0.180) |
|  | Experiment 3 (Amazon Mturk) ** | 0.339 (0.262) | N/A |
| Trials required to reach 0.9 test-retest reliability | Experiment 1 (College students), Visit 1 | 7 | Failed to reach 0.9 even after 42 trials |
|  | Experiment 1 (College students), Visit 2 | 6 | 39 |
|  | Experiment 2 (Patients w/ SUDs) | 11 | 27 |
|  | Experiment 3 (Amazon Mturk) ** | 11 | N/A |

**\*\* Note:** Except for Experiment 3 (Amazon Mturk participants), all experiments used 42 trials per session. In the Amazon Mturk experiment, there were 20 trials per session.

Lastly, we examined the correlation between the two model parameters (*k*: discounting rate and *β*:inverse temperature rate) when we used ADO or SC. Typically a high correlation between model parameters is undesirable because these parameters may influence each other ^e.g.,45^ and such a high correlation may lead to unstable parameter estimates. This was not the case for ADO; we found non-significant or weak (Pearson) correlation between log(k) and 1/*β*. Note that we separately calculated correlation coefficients in the two sessions within each visit. In Experiments 1 and 2, all but one correlation were non-significant and correlation coefficients never exceeded 0.25. In Experiment 3, with its very large sample, the correlation coefficients between the two parameters were 0.062 (p = 0.091) and 0.094 (p = 0.011), respectively, across the two sessions.

In contrast, with SC, the correlations between the two measures were much stronger^45^ in comparison to ADO. In Experiment 1, the correlation coefficients between the two parameters were at least 0.470 and all were highly significant (p < 0.0003). In Experiment 2, correlation coefficients were 0.340 (p = 0.066) and 0.229 (p = 0.223). Note that only ADO was used in Experiment 3. We believe these results further demonstrate the superiority of ADO over SC given that ADO leads to reduced correlations between model parameters in comparison to SC.

## Discussion

In three different populations, we have demonstrated that ADO led to highly reliable, precise, and rapid measures of discounting rate. ADO outperformed the SC method in college students (Experiment 1) and in patients meeting DSM-5 criteria for SUDs (Experiment 2). It held up very well in a less restrictive testing environment with a broader sample of the population (Experiment 3). The results of this study are consistent with previous studies employing ADO^29,46^, showing improved precision and efficiency. This is the first study demonstrating the advantages of ADO-driven delay discounting in healthy controls and psychiatric/online populations. In addition, this is one of the first studies that rigorously tested the precision of a latent measure (i.e., discounting rate) of a cognitive task. Such information is invaluable when evaluating methods and when making inferences from parameter estimates, as high precision can increase confidence. The SC method is an impressive heuristic method that delivers such good TRR (close to 0.90 in our study) that there is little room for improvement. Nevertheless, ADO is able to squeeze out additional information to increase reliability further. Where ADO does excel relative to the SC method is in precision and efficiency. The model-guided Bayesian inference that underlies ADO is responsible for this improvement. Unlike the SC method, which follows a simple rule of increasing or decreasing the value of a stimulus, ADO has no such constraint, choosing the stimulus that is expected to be most informative on the next trial. Trial after trial, this flexibility pays significant dividends in precision and efficiency, as the results of the three experiments show. **Figures S9**-**S10** illustrate how ADO and SC methods select upcoming designs (i.e., experimental parameters) in a representative participant (**Figure S9** at visit 1 and **Figure S10** at visit 2). In many cases, ADO quickly navigates to a small region of the design space. Interestingly, the selected region is often very consistent across multiple sessions and visits. SC shows a similar pattern but the stimulus-choice rule (the staircase algorithm) constrains to choose among pre-determined neighboring designs, making it less flexible, and thus requiring more trials, than ADO.

In all fairness, the above benefits of ADO also come with costs. For example, trials that are most *informative* can be ones that are also difficult for the participant^47^. Repeated presentation of difficult trials can frustrate and fatigue participants. Another issue is that for participants who respond consistently, the algorithm will quickly narrow (fewer than 10 trials) to a small region of the design space and present the same trials repeatedly with the goal of increasing precision even further. It is therefore important to implement measures that mitigate such behavior. We did so in the present study by inserting easy trials among difficult ones once the design space narrowed to a small number of options, keeping the total number of trials fixed. Another approach is to implement stopping criteria, such as ending the experiment once parameter estimation stabilizes, which would result in participants receiving different numbers of trials.

Both ADO and SC methods are different versions of a delay discounting task, and as such led to slightly different values of discounting rates: Correlations between *k* from ADO and *k* from the SC method ranged from 0.733 to 0.903 in Experiment 1 (four comparisons). It is unfortunate that no one truly knows the underlying discounting rate, but the fact that the association is not consistently high should not be surprising as it is true for any measures of human performance (e.g., measures of IQ, depression, or anxiety). Also, as mentioned above, ADO is more flexible than the SC method in the design choices selected from trial to trial. While the SC method is constrained to choosing among a few neighboring designs, there are no such limitations on ADO. This difference in flexibility will impact the final parameter estimate, especially in a short experiment.

There are a few reasons why we prefer the ADO approach. First, the reason that the correlation between ADO and SC was not always high (>.9) is likely due to noise, not the tasks, which are indistinguishable except for the sequencing of experimental parameters (e.g., reward amount and delay pairs) across trials. The greater precision of ADO compared to SC (3-5 times greater) suggests that SC is the larger source of this noise (see **Table 1** and **Figure S4**). Second, the reason for ADO’s greater precision is known, and lies in the ADO algorithm, which seeks to maximize information gain on each trial. There is a theoretically-motivated objective being achieved in the ADO approach that justifies stimulus choices trial after trial, whereas the SC approach is not as principled. More specifically, in the large sample theory of Bayesian inference, the posterior distribution is asymptotically normal, thus unimodal and symmetric, and also importantly, the posterior mean is optimal (‘consistent’ in a statistical sense), meaning that it converges to the underlying ground truth as the sample size increases^48^. Additionally, as shown earlier, two model parameters were statistically uncorrelated with ADO, which was not the case with SC. Together, the transparency of how ADO works along with its high reliability, precision, and parameter stability are strong reasons to prefer ADO. In summary, while we cannot say whether estimates using ADO are closest to individuals’ true internal states, ADO’s high consistency within and especially across visits (**Figure S5**) demonstrates a degree of trustworthiness.

While we believe that ADO is an exciting, promising method that offers the potential to advance the current state of the art in experimental design in characterizing mental functioning, we should mention a few major challenges and limitations in its practical implementation. One is the requirement that a computational/mathematical model of the experimental task be available. Also, the model should provide a good account (fit) of choice behavior. We believe the success of ADO in the delay discounting task is partly due to the availability of a reasonably good and simple hyperbolic model with just two free parameters. However, while we demonstrate the promise of an ADO method only in the area of delay discounting in this work, our methodology can be easily extended to other cognitive tasks that are of interest to researchers in psychology, decision neuroscience, psychiatry, and related fields. For example, we are currently applying ADO to tasks involving value-based or social decision making, including choice under risk and ambiguity^49^ and social interactions ^e.g., 50^. Preliminary results suggest that the superior reliability, precision, and efficiency of ADO compared to conventional methods generalize to other tasks. In addition, our recent work also demonstrates that ADO can be used to optimize the sequencing of stimuli and improve functional magnetic resonance imaging (fMRI) measurement^51^, which can reduce the cost of data acquisition and improve the quality of neuroimaging data.

Lastly, the mathematical details of ADO and its implementation in experimentation software can be a hurdle for researchers and clinicians. To reduce such barriers and allow even users with limited technical knowledge to use ADO in their research, we are developing user-friendly tools such as a Python-based package called *ADOpy*^52^ as well as web-based [www.mybrainstudy.com] and smartphone platforms.

In conclusion, the results of the current study suggest that machine-learning based tools such as ADO can improve the measurement of latent neurocognitive processes including delay discounting, and thereby assist in the development of assays for characterizing mental functioning and more generally advance measurement in the behavioral sciences and precision medicine in mental health.

## Methods

Reliability, precision, and efficiency were measured using the following quantitative definitions: Reliability was measured using the concordance correlation coefficient (CCC), which assesses the agreement between measurements collected at two points in time^44^. It is superior to the Pearson correlation coefficient, which assesses only association but not agreement. Precision was measured using within-subject variability quantified as the standard deviation of an individual parameter posterior distribution. Efficiency was quantified as the number of trials required to reach 0.9 TRR.

### Experiment 1 (college students)

#### Participants

Fifty-eight adult students at The Ohio State University (25 males and 33 females; age range 18-37 years; mean 19.0, SD 2.9 years) were recruited and received course credit for their participation. For all studies reported in this work, we used the following exclusion criterion: a participant is excluded from further analysis if the participant’ standard deviation (SD) of a parameter value (individual parameter posterior distribution) is two SD greater than the group mean. In other words, we excluded participants who seemingly made highly inconsistent choices during the task.

#### Delay discounting task

Each participant completed two sessions at each of the two visits. The two visits were separated by approximately one month (mean=28.3 days, SD=5.3 days). In each visit, a participant completed four delay discounting tasks: two ADO-based tasks and two SC-based tasks. Each ADO-based or SC-based task included 42 trials. The order of task completion (ADO then SC versus the reverse) was counterbalanced across participants.

In the traditional SC method, a participant initially made a choice between $400 now and $800 at seven different delays: one week, two weeks, one month, six months, one year, 3 years, and 10 years. Order of the delays was randomized for each participant. By adjusting the immediate amount, the choices were designed to estimate the participant’s indifference point for each delay. Specifically, the immediate amount was updated after each choice in increments totaling 50% of the preceding increment (beginning with $200), in a direction to make the unchosen option more subjectively valuable. For example, when presented with $400 now or $800 in 1 year, selecting the immediate option will lead to a choice between $200 now or $800 in 1 year ($200 increment). Then, choosing the immediate option once more will lead to a choice between $100 now or $800 in 1 year ($100 increment). If the later amount is then chosen, the next choice will be between $150 now or $800 in 1 year ($50 increment). This adjusting procedure ends after receiving choices for the subsequent $25 and $12.5 increments. See ^11,14^ for more examples of the procedure.

In the ADO method, the sooner delay and a later-larger reward were fixed as 0 day and $800. A later delay and a sooner reward were experimental parameters that were optimized on each trial. Based on the ADO framework and the participant’s choices so far, the most informative design (a later delay and a sooner reward) was selected on each trial. See prior publications^24,25^ and also **Figure 1** for technical details of the ADO framework.

#### Computational modeling

We applied ADO to the hyperbolic function, which has two parameters (*k*: discounting rate and *β*:inverse temperature rate). The hyperbolic function has the form *V = A / (1* + *kD)*, where an objective reward amount *A* after delay *D* is discounted to a subjected reward value *V* for an individual whose discounting rate is *k* (>0). In a typical delay discounting task, two options are presented on each trial: a smaller-sooner (SS) reward and a later-lager (LL) reward. The subjective values of the two options are modeled by the hyperbolic function. We used softmax (Luce’s choice rule) to translate subjective values into the choice probability on trial *t*:

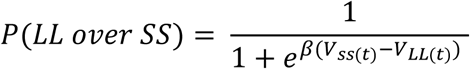

Where *V*_*SS*_ and *V*_*LL*_ are subjective values of the SS and LL options. To estimate the two parameters of the hyperbolic model in the SC method, we used the hBayesDM package^53^. The hBayesDM package (https://github.com/CCS-Lab/hBayesDM) offers hierarchical and non-hierarchical Bayesian analysis of various computational models and tasks using Stan software^54^. The hBayesDM function of the hyperbolic model for estimating a single participant’s data is *dd_hyperbolic_single*. Note that updating of our ADO framework is based on each participant’s data only. Thus, for fair comparisons between ADO and SC methods, we used an individual (non-hierarchical) Bayesian approach for analysis of data from the SC method. In ADO sessions, parameters (means and SDs of the parameter posterior distributions of the hyperbolic model) are automatically estimated on each trial. Note that estimation of discounting rate (*k*) was of primary interest in this project. We found that the TRR of the inverse temperature rate of the softmax function is much lower than that of *k* (discounting rate). We do not have a satisfying explanation and future studies are needed to investigate this issue. Estimates of the inverse temperature rate (a measure of response consistency or a degree of exploration/exploitation), *β*, are provided in the Supplemental Figures.

### Experiment 2 (patients meeting criteria for a substance use disorder)

#### Participants

Thirty-five individuals meeting DSM-5 criteria for a SUD and receiving treatment for addiction problems participated in the experiment (25 males and 10 females; age range 22-57 years; mean 35.8, SD 10.3 years). All participants were recruited through in-patient units at The Ohio State University Wexner Medical Center where they were receiving treatment for addiction. Trained graduate students and a study coordinator (Y.S.) used the Structured Clinical Interview for DSM-5 disorders (SCID-5) to assess diagnosis of a SUD. Final diagnostic determinations were made by W.-Y. A. on the basis of patients’ medical records and the SCID-5 interview. Exclusion criteria for all individuals included head trauma with loss of consciousness for over 5 minutes, a history of psychotic disorders, history of seizures or electroconvulsive therapy, and neurological disorders. Participants received gift cards for their participation (worth of $10/hr).

#### Delay discounting task and computational modeling

The task and methods for computational modeling in Experiment 2 were identical to those in Experiment 1. For a subset of participants in Experiment 2 (15 out of 35), the upper bound for discounting rate (*k*) during ADO was set as 0.1 for computing efficiency and we noted that some participants’ *k* values reached ceiling (=0.1). For the other participants (n=20), the upper bound was set to 1. We report results that are based on all 35 patients (**Figure 3A**) as well as results without participants whose *k* values reached the ceiling of 0.1 (**Figure S7**).

### Experiment 3 (large online sample)

#### Participants

Eight hundred and eight individuals through Amazon Mechanical Turk (MTurk; 353 males and 418 females (37 individuals declined to report their sex); age mean 35.0, SD 10.8 years) were recruited. They were required to reside in the United States and be at least 18 years of age, and received $10/hr for their participation. Out of 808 participants, 71 participants (8.78%) were excluded based on exclusion criteria (see Experiment 1).

#### Delay discounting task

Each participant completed two consecutive ADO-based tasks (sessions). The tasks were identical to the ADO version in Experiments 1 and 2 but consisted of just 20 trials per session (c.f., 42 trials per session in Experiments 1 and 2). There was no break between the two tasks, so participants experienced the experiment as a single session.

All participants received detailed information about the study protocol and gave written informed consent in accordance with the Institutional Review Board at The Ohio State University, OH, USA. All experiments were performed in accordance with relevant guidelines and regulations at The Ohio State University. All experimental protocols were approved by the Ohio State University Institutional Review Board.

## Acknowledgement

The research was supported by National Institute of Health Grant R01-MH093838 to M.A.P. and J.I.M, the Basic Science Research Program through the National Research Foundation (NRF) of Korea funded by the Ministry of Science, ICT, & Future Planning (NRF-2018R1C1B3007313 and NRF-2018R1A4A1025891) to W.-Y.A., the Institute for Information & Communications Technology Planning & Evaluation (IITP) grant funded by the Korea government (MSIT) (No. 2019-0-01367, BabyMind), and the Creative-Pioneering Researchers Program through Seoul National University to W.-Y.A. We thank Andrew Rogers and Zoey Butka for their assistance in data collection.

## Author contributions

W.-Y.A., J.M., and M.P. conceived and designed the experiments. Y.S., N.H., and H. H. collected data. W.-Y.A. performed the data analysis and drafted the paper. J.M. and M.P. co-wrote subsequent drafts with W.-Y.A. H.G. performed the data analysis. All authors (W.-Y.A., H.G., Y.S., N.H., H.H., J.T., J.M., and M.P.) contributed to writing the manuscript and approved the final version of the paper for submission.

