## Supplementary material for "Rapid, precise, and reliable measurement of delay discounting using a Bayesian learning algorithm"

**Figure S1**. Within-visit test-retest reliability of temporal discounting rates across two ADO (A-B) sessions and SC (C-D) sessions (Experiment 1, college students, with all 42 trials per session). Error bars on each point represent ±1 standard deviation within a participant. The line represents a linear regression line included to aid viewers in evaluating the linear trend in the data. Shaded regions represent ±1 standard deviation across participants. CCC: Concordance Correlation Coefficient.

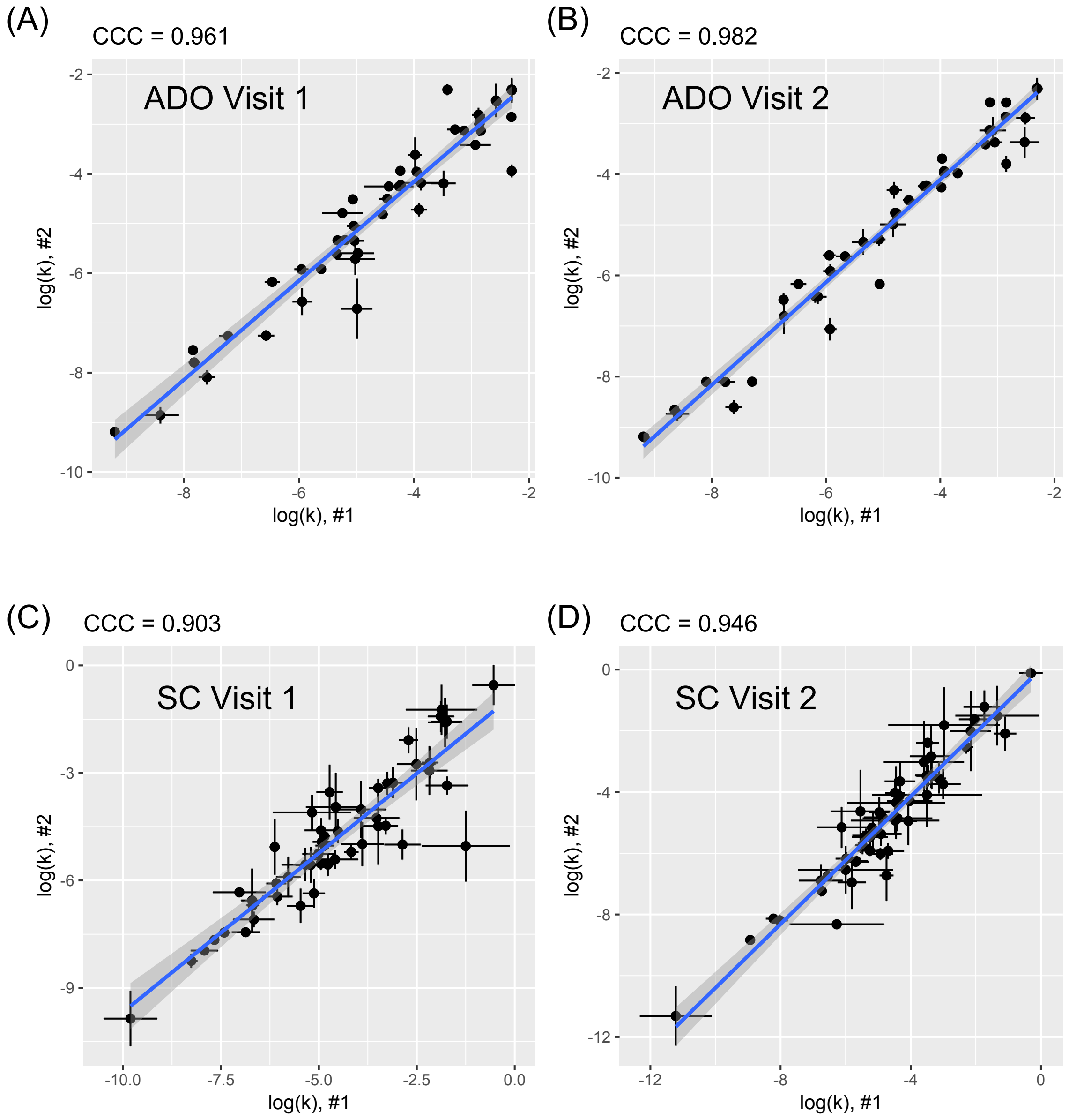

**Figure S2.** Within-visit test-retest reliability of temporal discounting rates (log(k), A & C) and inverse temperature parameters (β, B & D) among college students (Experiment 1) with ADO including outliers, which are indicated as red circles. See **Methods and Materials** for the description of outliers. (A & B) At visit 1, (C & D) At visit 2, which was separated by approximately one month from visit 1. Error bars on each point represent ±1 standard deviation within a participant. The line represents a linear regression line included to aid viewers in evaluating the linear trend in the data. Shaded regions represent ±1 standard deviation across participants. CCC: Concordance Correlation Coefficient.

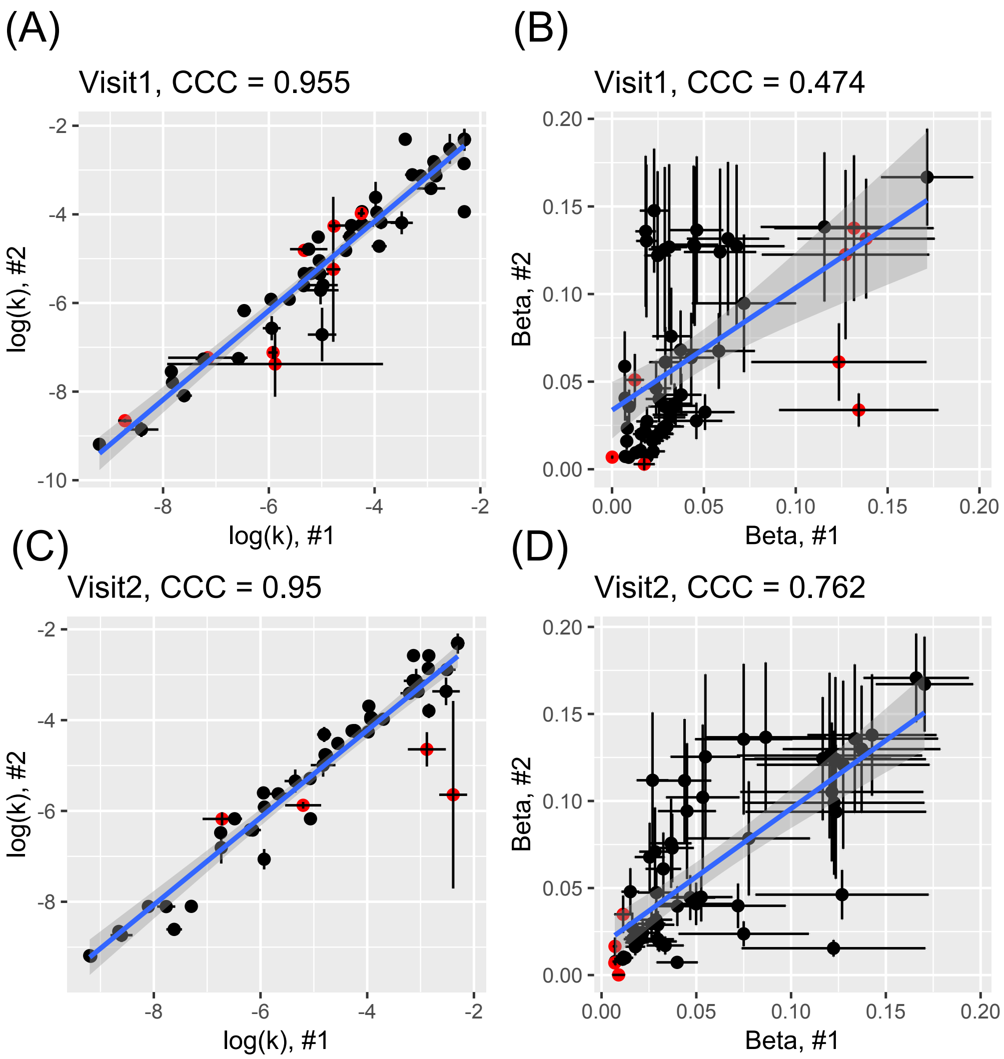

**Figure S3.** Within-visit test-retest reliability of temporal discounting rates (log(k), A & C) and inverse temperature parameters (β, B & D) among college students (Experiment 1) with the staircase method including outliers, which are indicated as red circles. See **Methods and Materials** for the description of outliers. (A & B) At visit 1, (C & D) At visit 2, which was separated by approximately one month from visit 1. Error bars on each point represent ±1 standard deviation within a participant. The line represents a linear regression line included to aid viewers in evaluating the linear trend in the data. Shaded regions represent ±1 standard deviation across participants. CCC: Concordance Correlation Coefficient.

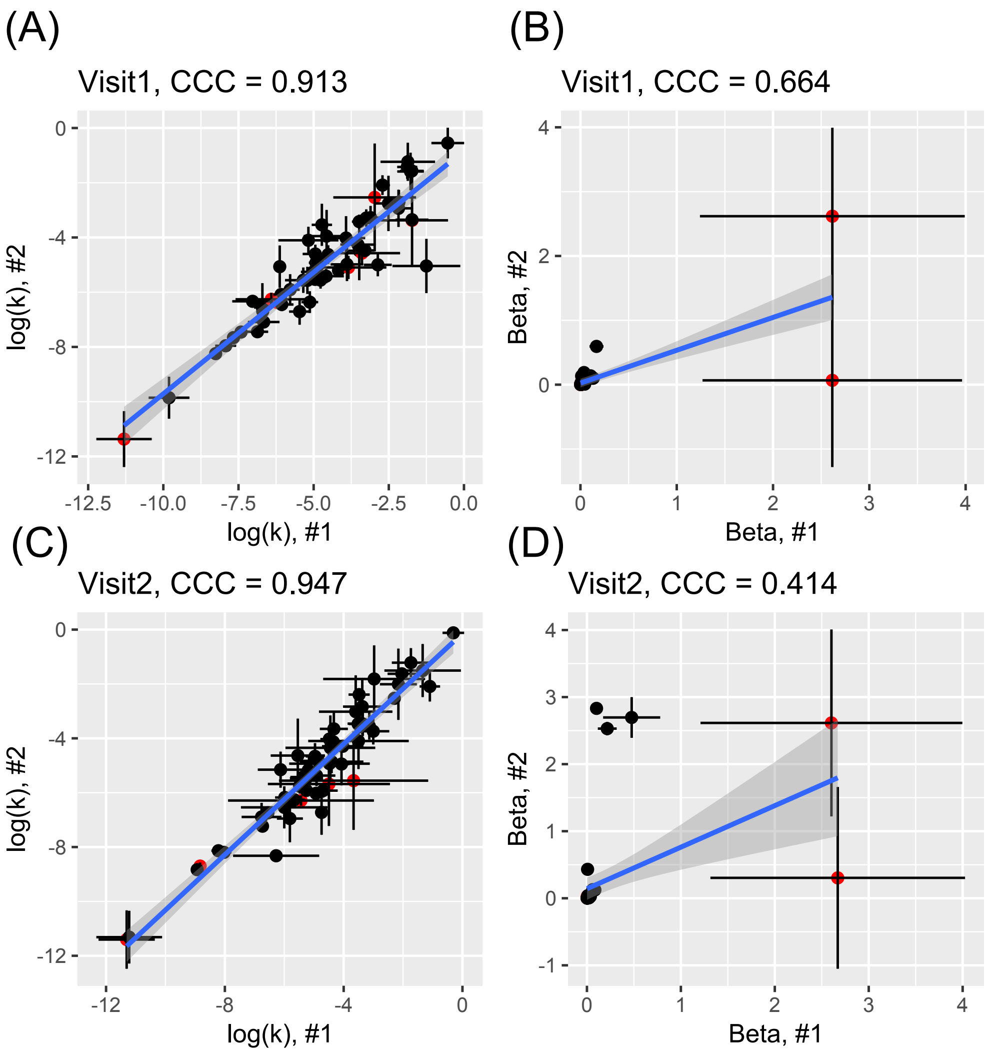

**Figure S4**. Comparison of ADO and staircase methods with respect to their precision of parameter estimates (Experiment 1, college students). As a measure of precision, we used the standard deviation (SD) of an individual parameter posterior (natural log of the posterior distribution of the temporal discounting rate (k)). Thus, the smaller SD is, the greater its precision is. (A) At visit 1 (B) At visit 2, which was separated by approximately one month from visit 1. The SDs are 0.122 (visit 1) and 0.098 (visit 2) for ADO, and 0.413 (visit 1) and 0.537 (visit 2) for SC.

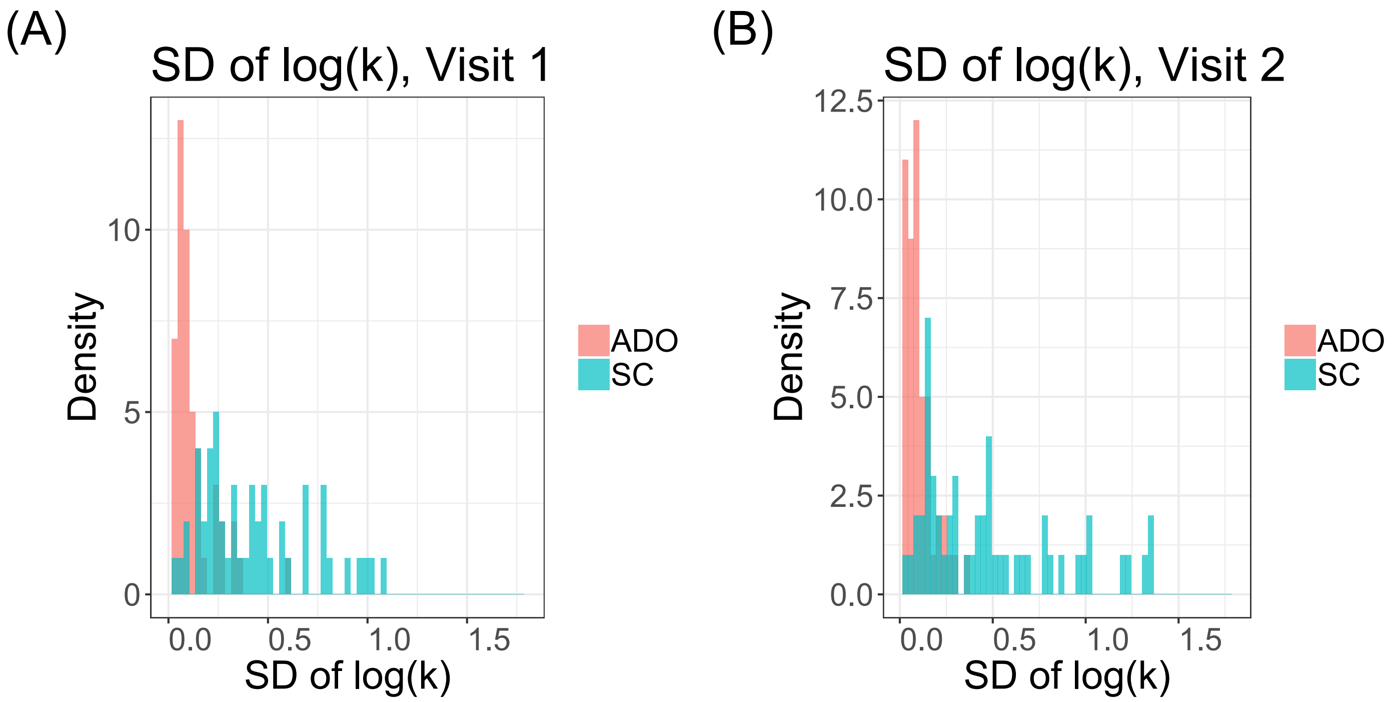

**Figure S5**. Comparison of between-visit efficiency of the ADO (A) and the staircase (B) methods (Experiment 1, college students). Between-visit efficiency is measured as the cumulative test-retest reliability in each trial (ADO) or every third trial (staircase) across two visits. The two visits were separate by approximately one month. Each line represents a different test-retest reliability comparison across the two visits (1. visit1-session1 vs visit2-session1 (red); 2. visit1-session1 vs visit2-session2 (green); 3. visit1-session2 vs visit2-session1 (cyan); 4. visit1-session2 vs visit2-session2 (purple)).

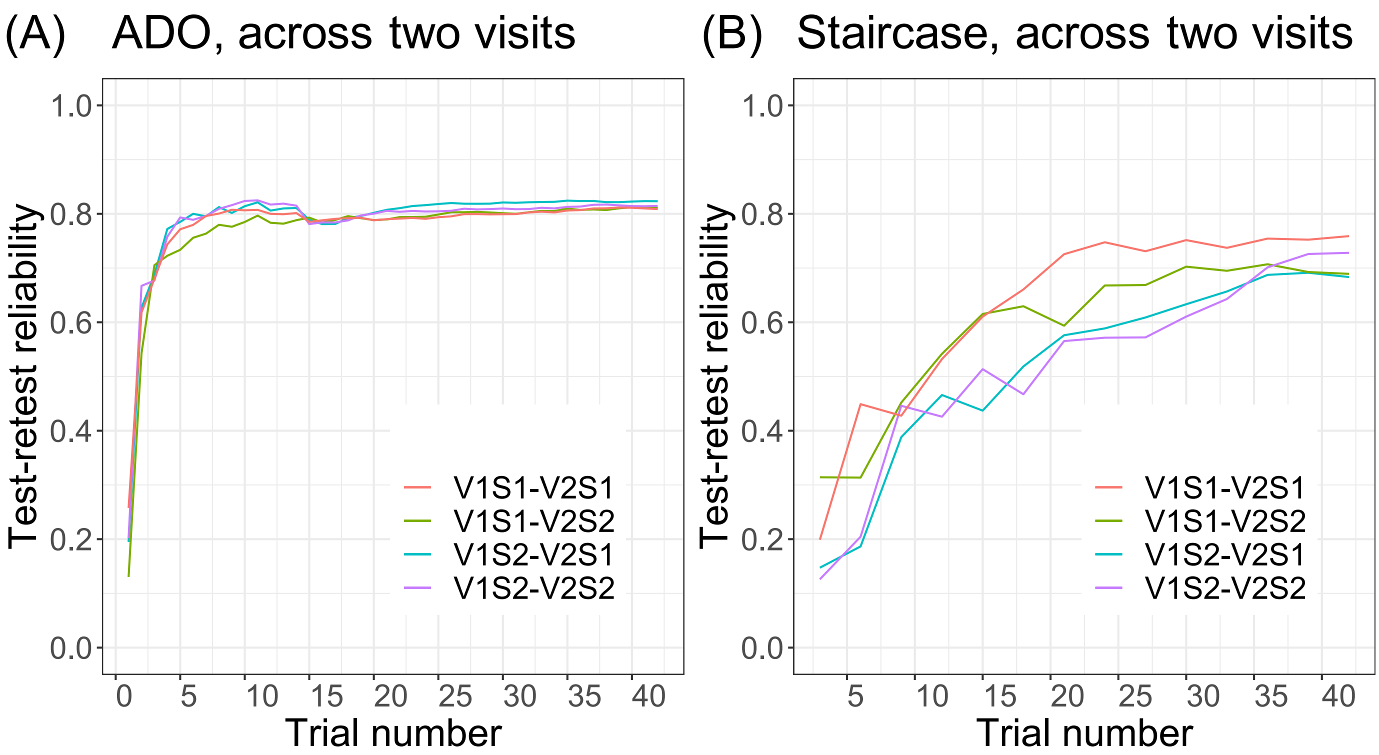

**Figure S6.** Test-retest reliability of discounting rates (log(k), A & C) and inverse temperature parameters (β, B & D) among patients with substance use disorders (Experiment 2) with ADO and staircase methods including outliers, which are indicated as red circles. See **Methods and Materials** for the description of outliers. (A & B) With ADO, (C & D) With the staircase method. Error bars on each point represent ±1 standard deviation within a participant. The line represents a linear regression line included to aid viewers in evaluating the linear trend in the data. Shaded regions represent ±1 standard deviation across participants. CCC: Concordance Correlation Coefficient.

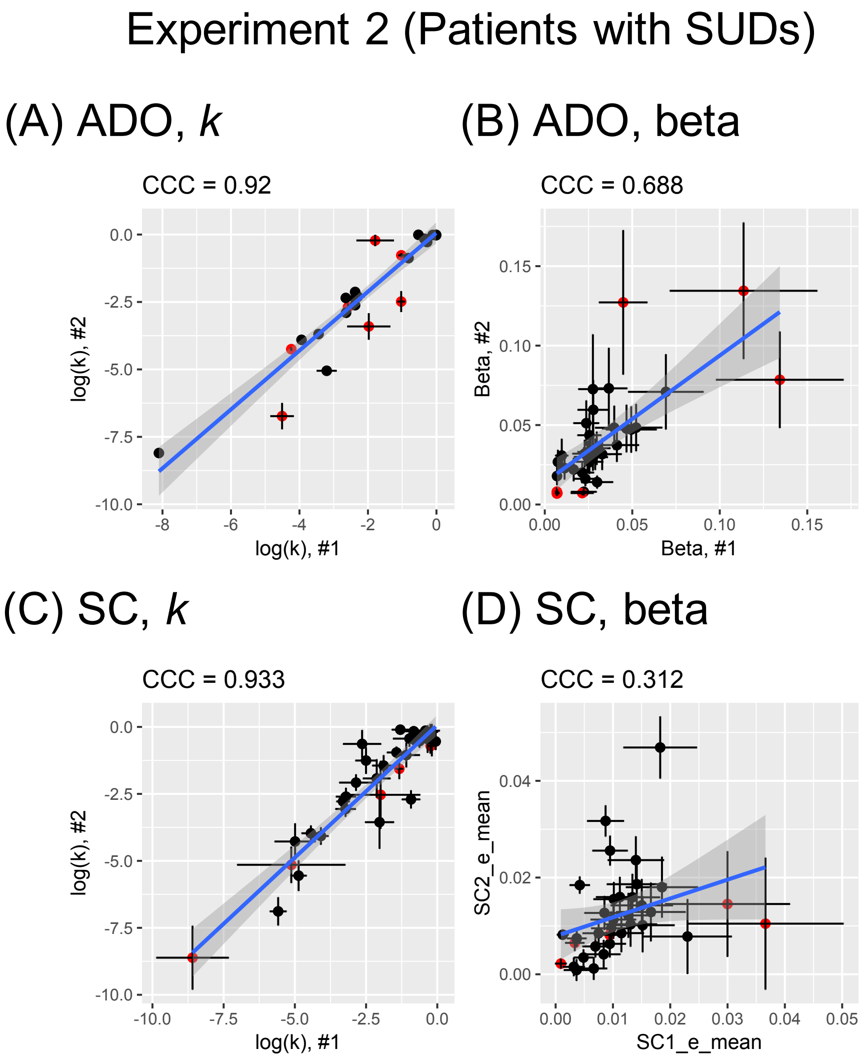

**Figure S7.** Reliability and efficiency of the staircase method in Experiment 2 (N=15 patients with substance use disorders (SUDs)) with ADO. Out of 35, 14 patients whose *k* values reached the upper bound (=0.1) were first excluded and 6 additional patients were excluded with the 2SD rule. (A) Test-retest reliability with all 42 trials per session. The line represents a linear regression line included to aid viewers in evaluating the linear trend in the data. Shaded regions represent ±1 standard deviation across participants. (B) Efficiency metric using cumulative test-retest reliability across trials. Shaded regions represent ±1 standard deviation across participants. CCC: Concordance Correlation Coefficient.

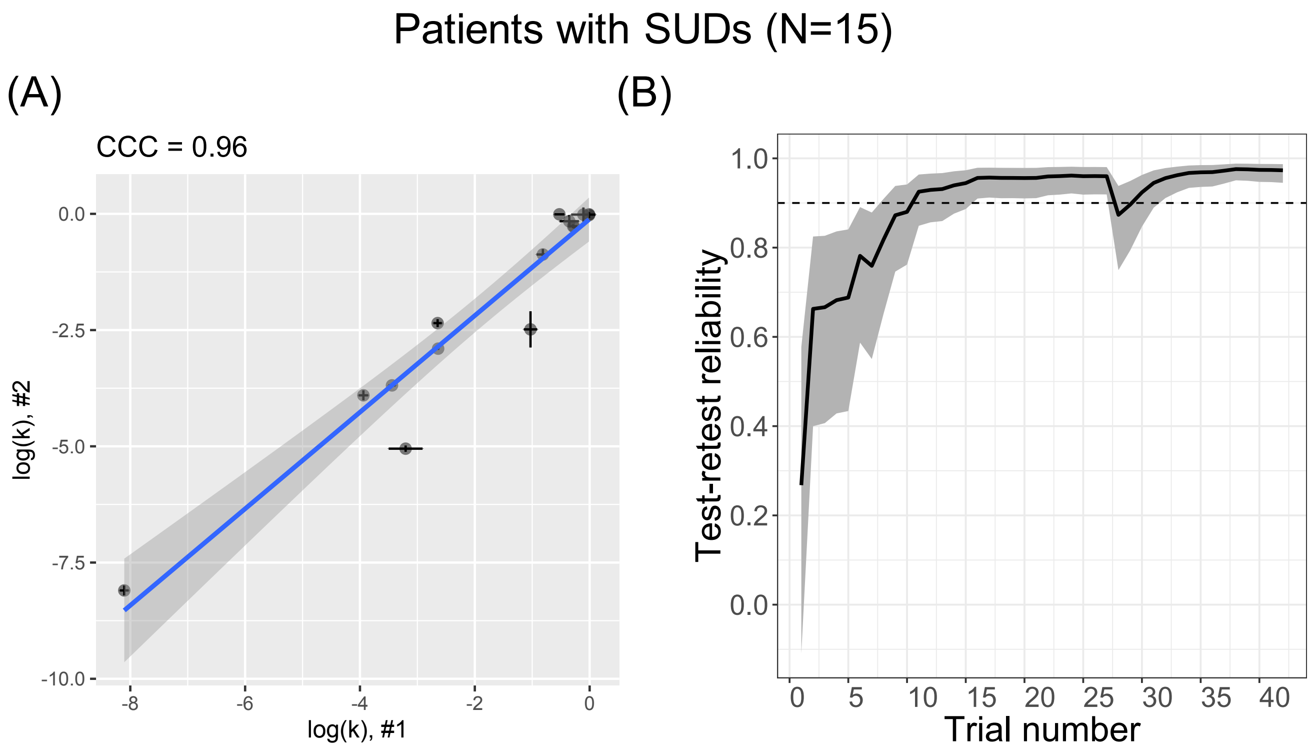

**Figure S8.** Test-retest reliability of discounting rates (log(k)) and inverse temperature parameters (β) among large online Amazon MTurk participants (Experiment 3) with ADO including outliers, which are indicated as red circles. See **Methods and Materials** for the description of outliers. Error bars on each point represent ±1 standard deviation within a participant. Shaded regions represent ±1 standard deviation across participants. Shaded regions represent ±1 standard deviation across participants. CCC: Concordance Correlation Coefficient.

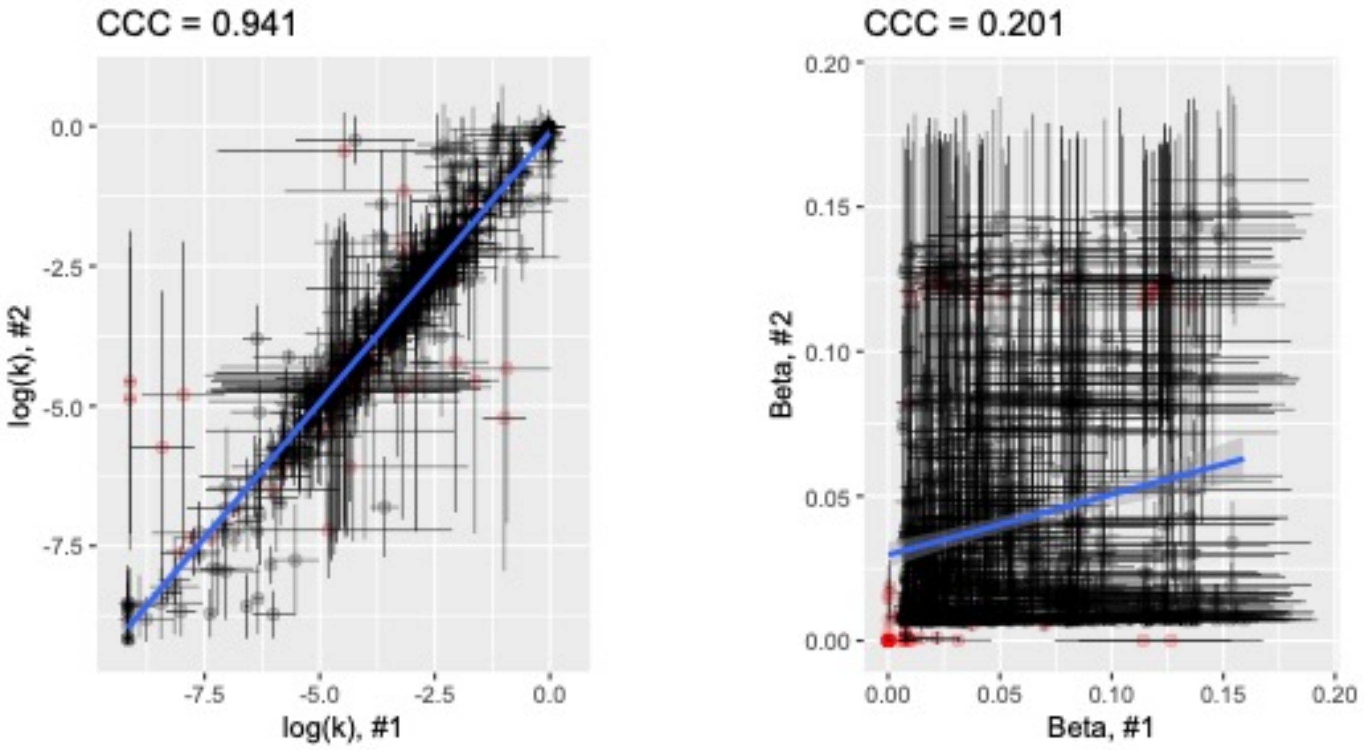

**Figure S9.** Trajectories of design space selection over trials with ADO and Staircase (SC) for a representative participant in Experiment 1 and visit 1 (see **Figure S10** for the trajectories of design space selection in visit 2 of the same participant). Design space refers to two experimental parameters: (1) delays for larger and later (LL) rewards and (2) amount for smaller and sooner (SS) rewards. Color bars indicate trial numbers (1 through 42 represented by a transition from blue to red).

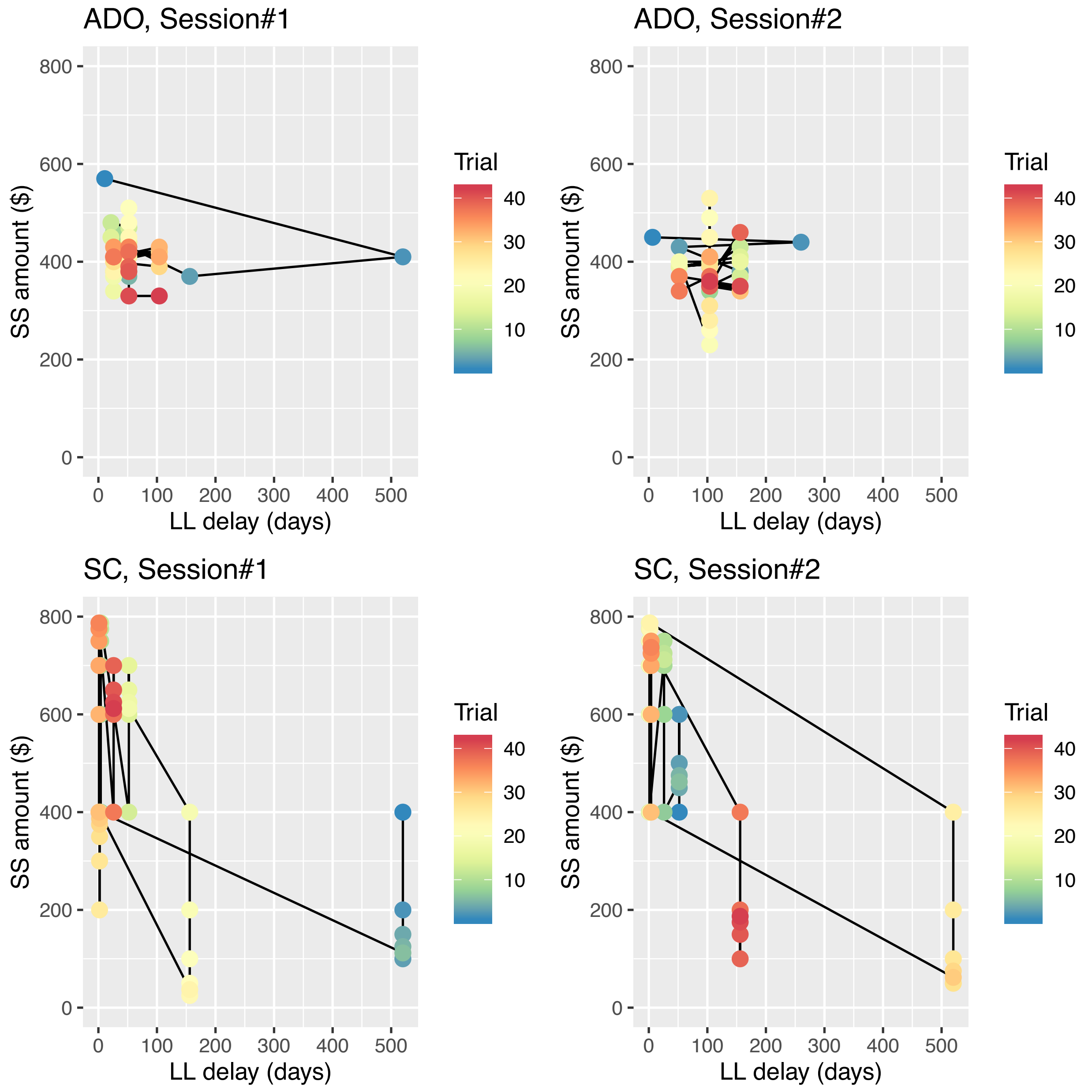

**Figure S10.** Trajectories of design space selection over trials with ADO and Staircase (SC) for a representative participant in Experiment 1 and visit 2 (see **Figure S9** for the trajectories of design space selection in visit 1 of the same participant). Design space refers to two experimental parameters: (1) delays for larger and later (LL) rewards and (2) amount for smaller and sooner (SS) rewards. Color bars indicate trial numbers (1 through 42 represented by transitions from blue to red).

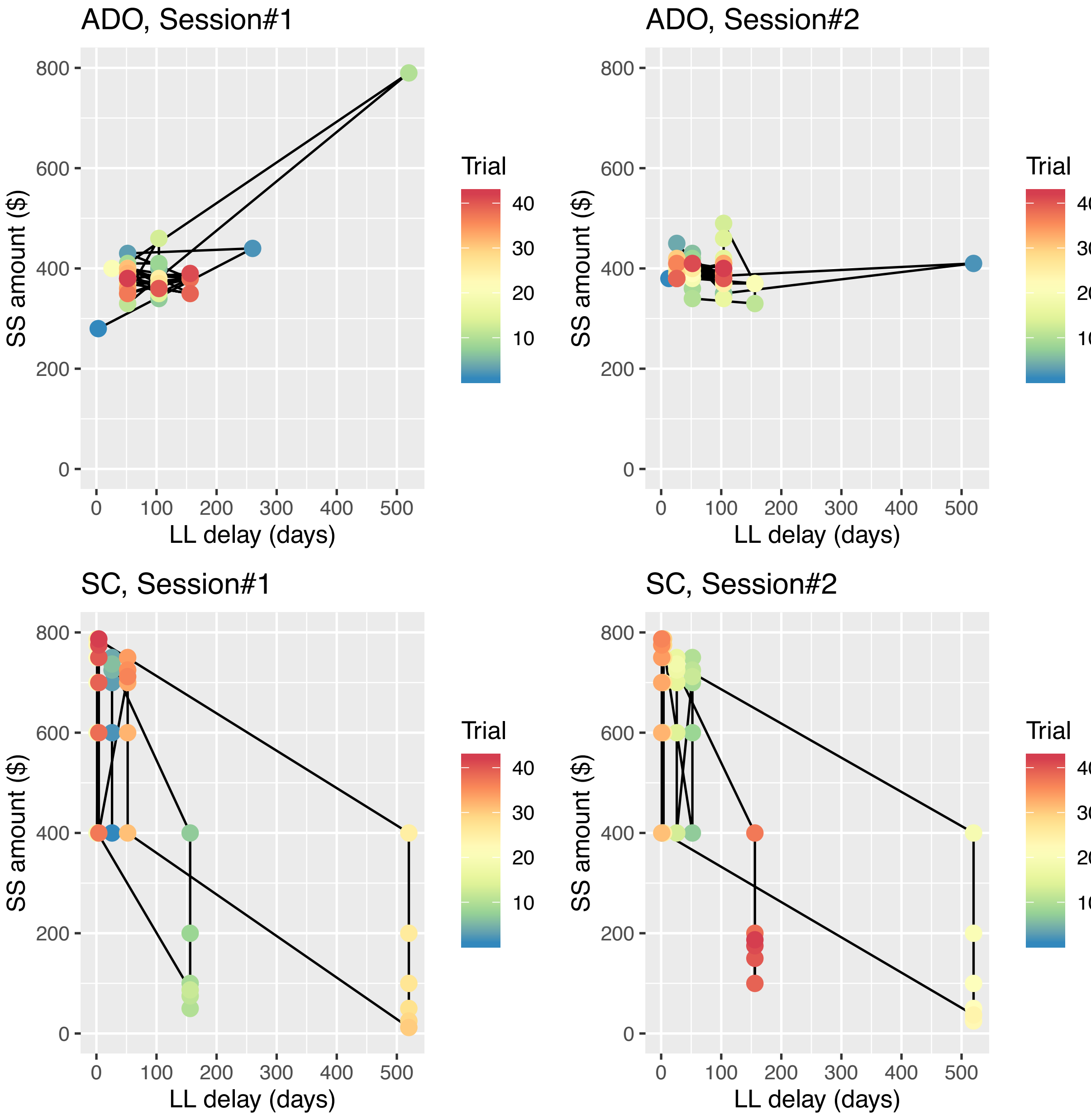
